# Comparison of gastric inflammation and metaplasia induced by *Helicobacter pylori* or *Helicobacter felis* colonization in mice

**DOI:** 10.1101/2023.12.22.573128

**Authors:** Sara R. Druffner, Shrinidhi Venkateshwaraprabu, Stuti Khadka, Benjamin C. Duncan, Maeve T. Morris, Emel Sen-Kilic, Fredrick H. Damron, George W. Liechti, Jonathan T. Busada

**Author notes:** **Corresponding authors:** Jonathan T. Busada, WVU School of Medicine, Microbiology, Immunology and Cell Biology, 64 Medical Center Drive, P.O. Box 9177, Morgantown, WV 26506., Phone: -4621. Joint first author (Sara R. Druffner, Shrinidhi Venkateshwaraprabu). **Disclosures:** The authors have declared that no conflict of interest exists.

## Abstract

**Background:** Gastric cancer is the fifth most diagnosed cancer in the world. Infection by the bacteria *Helicobacter pylori* (HP) is associated with approximately 75% of gastric cancer cases. HP infection induces chronic gastric inflammation, damaging the stomach and fostering carcinogenesis. Most mechanistic studies on *Helicobacter-*induced gastric cancer initiation are performed in mice and utilize either mouse-adapted strains of HP or the natural mouse pathogen *Helicobacter felis* (HF). Each of these infection models is associated with strengths and weaknesses. Here, we identified the differences in immunogenicity and gastric pathological changes associated with HP and HF infection in mice.

**Material and Methods:** PMSS1 HP strain or with the CS1 HF strain were co-cultured with mouse peritoneal macrophages to assess their immunostimulatory effects. C57BL/6J mice were infected with HP or HF, and gastric inflammation, atrophy, and metaplasia development were assessed 2 months post-infection.

**Results:** HP and HF induced similar cytokine production from cultured mouse peritoneal macrophages. HP-infected mice caused modest inflammation within both the gastric corpus and antrum and did not induce significant atrophy within the gastric corpus. In contrast, HF induced significant inflammation throughout the gastric corpus and antrum. Moreover, HF infection was associated with significant atrophy of the chief and parietal cell compartments and induced expression of pyloric metaplasia markers.

**Conclusions:** HP is poorly immunogenic compared to HF. HF induces dramatic CD4+ T cell activation, which is associated with increased gastric cancer risk in humans. Thus, HP studies in mice are better suited for studies on colonization, while HF is more strongly suited for pathogenesis and cancer initiation studies.

## Introduction

Gastric cancer is the 5th most common cancer and the 4th leading cause of cancer deaths worldwide ^(1)^. *Helicobacter pylori* (HP) infection is the primary risk factor for gastric cancer, contributing to 75% of the global gastric cancer burden ^(2)^. HP infection is extremely prevalent, infecting approximately 50% of the world’s population and 1-3% of infected individuals will develop cancer ^(3)^. Stomach infection by HP induces chronic inflammation of the gastric mucosa that in turn drives widespread epithelial damage and metaplasia’s, eventually resulting in adenocarcinoma ^(2)^. Gastric inflammation is necessary and sufficient for gastric cancer initiation. Sterile inflammation drives atrophic gastritis and metaplasia development, and mouse models with gastric-specific overexpression of the proinflammatory cytokines IL1B ^(4)^ or INFG ^(5)^ and mouse models of autoimmune gastritis develop gastric tumors in the absence of *Helicobacter* infection. Similarly, polymorphisms resulting in higher levels of the proinflammatory cytokines IL1B or TNF and lower levels of the anti-inflammatory cytokine IL10 are associated with increases in HP-related gastric cancer ^(6, 7)^. In contrast, multiple immune-deficient mouse models are protected from the gastric pathologies associated with *Helicobacter* infection ^(8-10)^. While it is well established that chronic inflammation modifies gastric cancer risk, the inflammatory phenotypes that promote cancer initiation remain poorly defined.

The vast majority of mechanistic studies are performed in mouse models. However, studies in mice pose specific challenges. HP is not a natural mouse pathogen, and clinical isolates typically require serial passage through mice to enhance mouse colonization ^(11, 12)^. Mouse-adapted HP strains, such as the commonly used SS1 strain, have lost expression of several virulence factors that are imported for causing disease in humans, and they rarely cause severe disease in mice ^(13)^. CagA is the best-described example of this loss of virulence factors. The CagA pathogenicity island is the best-known correlate of cancer risk. Encoding a type IV secretion system (T4SS) and CagA oncoprotein, this system injects CagA and other bacterial components into host epithelial cells, exacerbating inflammation and inducing host cell mutations ^(14)^. In the SS1 strain, CagA translocation to host epithelial cells is lost due to rearrangements in CagY ^(15)^. While the pre-mouse adapted SS1 strain (PMSS1) maintains CagA expression and T4SS function *in vitro*, CagA translocation is lost after 3-4 months of mouse colonization ^(16, 17)^

*Helicobacter felis* (HF) is a close relative to HP that was originally isolated from the gastric mucosa of a cat ^(18)^. The HF genome shares more than 60% homology with HP and encodes many orthologs of HP essential genes that are required for gastric colonization ^(19)^. HF readily colonizes mice and quickly triggers massive gastric inflammation and atrophic gastritis within 2 months post-infection ^(20)^. Long-term infection over 12-24 months induces dysplasia and occasionally non-invasive adenocarcinoma ^(21)^. While HF does colonize the human stomach, it is not a gastric cancer risk factor ^(22, 23)^. While the CagA pathogenicity island is absent in HF ^(19)^, its robust induction of gastric inflammation makes it well-suited as a model of HP pathogenesis in mice.

Both HP and HF are widely used to study gastric cancer initiation, but the strengths and weaknesses of each bacteria species may not be readily apparent to researchers. Typically, studies have elected to use one species or the other, and few have directly compared either the bacteria to one another or their effects on the host ^(24, 25)^. The goal of this study was to directly compare the effects of HP and HF on immune cell activation in vitro and their effects on gastric inflammation, atrophy, and metaplasia development in mice. We infected C57BL/6J mice with the HP PMSS1 strain, the most widely used CagA+ strain, and the HF CS1 strain. Mice were assessed 2 months post-infection to ensure that HP still maintained CagA function and as this represents a convenient timeframe for investigating the effects of chronic inflammation on the gastric epithelium. Our findings indicate that both bacterial species exhibit a similar ability to stimulate macrophages in vitro but that HF induces significantly more intense and widespread gastric inflammation and atrophy in mice. While both bacterial strains have specific utility, this study aims to serve as a resource to help researchers choose the bacterial model that best suits their experimental goals.

## Materials and Methods

### Animal Care and Treatment

All mouse studies were performed with approval by the West Virginia University Animal Care and Use Committee. C57BL/6J mice were purchased from the Jackson Laboratories. Mice were administered standard chow and water *ab libitum* and maintained in a temperature-and humidity-controlled room with standard 12-hour light/dark cycles. 8-week-old mice were mock infected or infected with HP or HF by oral gavage. Mock mice received 500μL sterile brucella broth, and infected mice were inoculated with 500 μL of brucella broth containing 10^9^ CFU of the respective bacteria two times 24 hours apart. For the HP infection studies, both male and female mice were used. While for the HF infection studies, only female mice were used as there are significant sex differences in the response to HF. For peritoneal macrophage isolation, mice received a single IP injection of 1mL sterile Brewers Thioglycollate media (Sigma-Aldrich). After 4 days peritoneal lavage was plated and nonadherent cells were removed by washing with prewarmed 1xPBS.

### Bacterial Preparation

HF (ATCC 49179) was grown on tryptic soy agar plates (BD Biosciences) with 5% defibrinated sheep blood (Hemostat Labs) and 10 mg/mL vancomycin (Alfa Aesar) under microaerophilic conditions) 5% O2 and 10% CO2) at 37°C for 2 days. HF was then harvested and transferred to Brucella broth (Research Products International) containing 5% fetal bovine serum (R&D Systems) and 10 mg/mL vancomycin and grown overnight at 37°C under microaerophilic conditions with agitation. Bacteria were centrifuged and resuspended in fresh Brucella broth without antibiotics before spectrophotometry and mouse infection.

*H pylori* PMSS1^(26)^ (a gift from Manuel Amieva, Stanford University) was inoculated in Brucella broth containing 10% fetal bovine serum and 10 mg/mL vancomycin and grown overnight at 37°C under microaerophilic conditions with agitation. Bacteria were centrifuged and resuspended in fresh Brucella broth without antibiotics before spectrophotometry and mouse infection.

### Tissue Preparation

Mice were euthanized 2 months post-inoculation. Stomachs were removed and opened along the greater curvature and washed in phosphate-buffered saline to remove gastric contents. One side of the stomach was fixed overnight in 4% paraformaldehyde at 4°C, and cut into strips. Strips were either cryopreserved in 30% sucrose and embedded in optimal cutting temperature (OCT) media or transferred into 70% ethanol and submitted to the West Virginia University histology core for routine processing, embedding, sectioning, and H&E staining. For histology, strips were taken from the corpus greater curvature. A 2mm biopsy was removed from the other side of the corpus greater curvature for RNA isolation and immediately snap-frozen in liquid nitrogen. The remained of the corpus was disassociated into a single cell suspension for flow cytometry as described below.

### Histology

Immunostaining was performed using standard methods. Briefly, 5-μm stomach cryosections were incubated with anti H+/K+ ATPase antibodies (clone 1H9, MBL Life Science), MIST1 (Cell Signaling Technologies), CD45 (clone 104; Biolegend) or CD44v9 (Cosmo Bio) for 1 hour at room temperature or overnight at 4°C. Sections were incubated in secondary antibodies for 1 hour at room temperature. Fluorescence-conjugated *Griffonia simplicifolia* lectin (GSII; ThermoFisher) was added with secondary antibodies where indicated. Sections were mounted with Vectastain mounting media containing 4’,6-diamidino-2-phenylindole (Vector Laboratories). Images were obtained using a Zeiss 710 confocal laser-scanning microscope (Carl-Zeiss) and running Zen Black (Carl-Zeiss) imaging software.

### RNA Isolation and qRT-PCR

For gastric tissue RNA was extracted in TRIzol (Thermo Fisher Scientific) and precipitated from the aqueous phase using 1.5 volumes of 100% ethanol. The mixture was transferred to an RNA isolation column (Omega Bio-Tek) and the remaining steps were followed according to the manufacturer’s recommendations. For peritoneal macrophages, RNA was isolated using the MicroElute Total RNA kit (Omega Bio-Tek). RNA was treated with RNase-free DNase I (Omega Bio-Tek) as part of the isolation procedure. Reverse transcription followed by qPCR was performed in the same reaction using the Universal Probes One-Step PCR kit (Bio-Rad Laboratories) and the TaqMan primers/probe mixtures (all from Thermo Fisher Scientific): *Ppib* Mm00478295_m1), *Wfdc2* (Mm00509434_m1), *Il13* (Mm00434204_m1), *Ifng* (Mm01168134_m1), *Cftr* (Mm00445197_m1), *Tnf* (Mm00443258_m1), and *Il1b* (Mm00434228_m1). Relative gene expression was normalized to *Ppib*.

### Flow Cytometry

Corpus tissue from euthanized mice was washed in Hanks Balanced Salt Solution without Ca^2+^ or Mg^2+^ containing 5 mM HEPES, 5 mM EDTA, and 5% FBS at 37°C for 20 minutes. Tissue was then washed in Hanks Balanced Salt Solution with Ca^2+^ or Mg^2+^ briefly and then digested in 1mg/ml collagenase (Worthington) for 30 minutes at 37°C. After digestion, the tissue fragments were pushed through a 100 μM strainer and then rinsed through a 40 μM strainer before debris was removed through an Optiprep (Serumwerk) density gradient. Fc receptors were blocked with TruStain (Biolegend) and then stained with antibodies for 20 minutes on ice. The following antibodies were used: CD45.2 (clone 104), CD3e (clone 145-2C11), B220 (clone RA3-6B2), CD4 (clone GK1.5), CD8a (clone 53-6.7), CD11b (clone M1/70), MHCII (clone M5/114.15.2) Ly6g (clone 1A8), F4/80 (clone BM8), and SiglecF (E50-2440). Actinomycin D (A1310, Invitrogen) was used to label dead cells. Cells were analyzed on a Cytek Aurora spectral flow cytometer (Cytek Biosciences). Flow cytometry analysis was performed using Cytobank (Beckman Coulter).

### Peritoneal Macrophage Treatment and Cytokine Array

Peritoneal macrophages were treated with vehicle (sterile brucella broth) or a 1:2 ratio of macrophages to HP or HF. Cells were washed and collected 3 hours post treatment for RNA isolation. Media was collected 24 hours post treatment for cytokine analysis. Cytokine were assessed using the Mouse C3 cytokine array (RayBiotech) following the manufacturers protocol. The arrays were image with an iBright 1500 (Thermofisher Scientific) and dot density was measured using the on-board analysis software. Cytokines were considered expressed if their pixel density was 1.5 times higher than the negative control dots printed on the cytokine array. A heatmap was generated using the Morpheus heat map tool (Broad Institute).

### Statistical Analysis

All error bars are ± SD of the mean. The sample size for each experiment is indicated in the figure legends. Experiments were repeated a minimum of 2 times. Statistical analyses were performed using 1-way analysis of variance with the post hoc Tukey t-test when comparing 3 or more groups or by an unpaired t-test when comparing two groups. Statistical analysis was performed by GraphPad Prism 10 software. Statistical significance was set at *P* -. 0.05. Specific *P* values are listed in the figure legends.

## Results

### H. felis and H. pylori exhibit similar immunostimulatory effects in vitro

HP has evolved to avoid the host immune system ^(27)^. Mutations in HP LPS and flagellar proteins avoid binding of TLR4 and TLR5, respectively ^(24, 28)^. The enhanced immunogenicity of HF in vivo raises the possibility that HF is intrinsically more immunostimulatory than HP. We isolated thioglycolate-induced peritoneal macrophages from WT C57Bl6/J mice to examine the immunostimulatory effects of HP and HF. The cell cultures were challenged with a 2:1 cell ratio of HP or HF for three hours before assessment of proinflammatory cytokine expression by qRT-PCR. Challenge with either bacterium induced similar levels of the *Tnf*, while *Il6* and *Il1b* induction was significantly higher in HF-challenged macrophages (Figure 1A). Next, we used the same experimental system to measure bacterial induction of cytokines. Macrophages were co-cultured with either HF or HP for 24 hours, and the cell culture media was assessed by a mouse 62 cytokine dot plot array (Table 1). For this assay, a cytokine was considered detectable if the densitometry was 1.5 fold above the negative control background. Twenty-two cytokines were detected in the tissue culture media. Several cytokines were dramatically induced by bacterial co-culture, but surprisingly, there were no notable differences between the cytokine expression profiles of HP or HF-stimulated macrophages (Figure 1 B-C). These data demonstrate that HP and HF have a similar propensity to stimulate macrophages in vitro, suggesting that both strains are similarly immunogenic.

**Figure 1.**
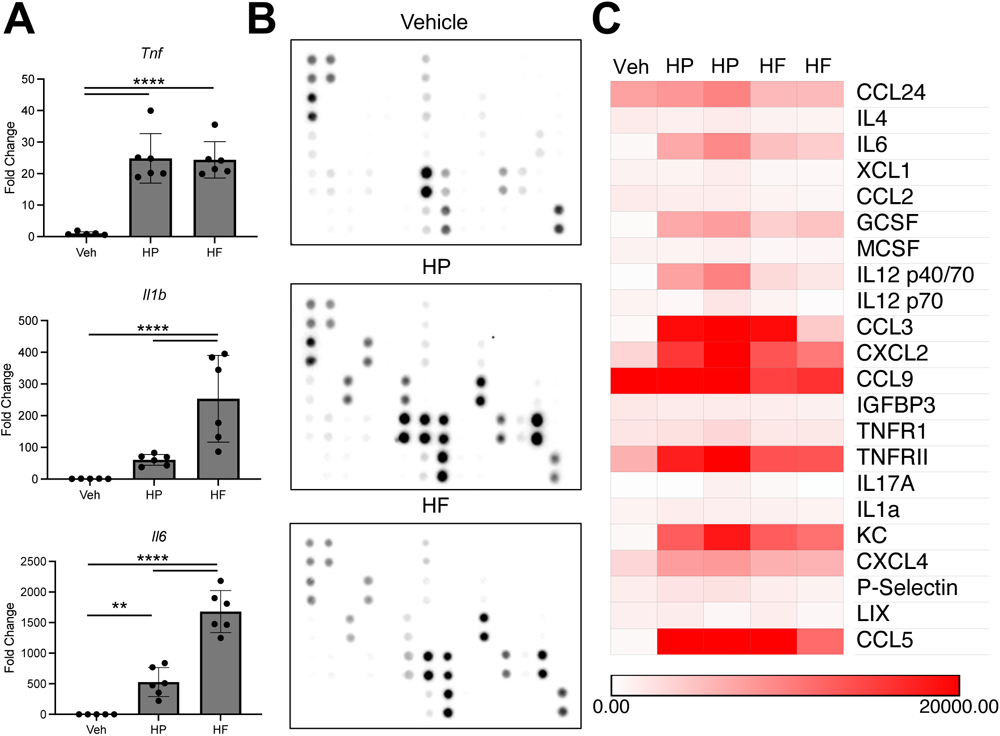
*H. pylori* and *H. felis* elicit similar activation of macrophages *in vitro.* **(A)** Quantitative RT-PCR of the indicated genes using RNA isolated from cultured peritoneal macrophages 3 hours after mock stimulation or stimulation with at 1:1 ratio of *H. pylori* or *H. felis.* n≥5. **p≤0.01; ****p≤0.0001. (**B**) Representative dot plot cytokine arrays. Arrays were probed with cell culture media from peritoneal macrophages stimulated for 24 hours with a 1:1 ratio of *H. pylori* or *H. felis.* Dot labels are found in Table 1. (**C**) Heatmap of the cytokine arrays shown in (B). Cytokines were considered expressed if the dot density was 1.5 fold above the background. n≥3.

**Table 1:** Cytokine array dot plot key.

|  | A | B | C | D | E | F | G | H | I | J | K | L | M | N |
| --- | --- | --- | --- | --- | --- | --- | --- | --- | --- | --- | --- | --- | --- | --- |
| 1 | Pos | Pos | Neg | Neg | Blank | AXL | CXCL13 | TNFSF8 | TNFRSF8 | TNFRSF5 | CRG2 | CCL27 | CXCL16 | CCL11 |
| 2 |  |  |  |  |  |  |  |  |  |  |  |  |  |  |
| 3 | CCL24 | TNFSF6 | CX3CL1 | GCSF | GMCSF | IFNγ | IGFBP3 | IGFBP5 | IGFBP6 | IL1α | IL1β | IL2 | IL3 | IL3Rβ |
| 4 |  |  |  |  |  |  |  |  |  |  |  |  |  |  |
| 5 | IL4 | IL5 | IL6 | IL9 | IL10 | IL12<br>p40/p70 | IL12 p70 | IL13 | IL17A | CXCL1 | Leptin R | Leptin | LIX | CD62L |
| 6 |  |  |  |  |  |  |  |  |  |  |  |  |  |  |
| 7 | XCL1 | CCL2 | MCP5 | MCSF | CXCL9 | CCL3 | MIP1γ | MIP2 | CCL19 | CCL20 | CXCL4 | P-Selectin | CCL5 | SCF |
| 8 |  |  |  |  |  |  |  |  |  |  |  |  |  |  |
| 9 | SDF1α | CCL17 | CCL1 | CCL25 | TIMP1 | TNFα | TNFRSF1A | TNFRSF1B | TPO | CD106 | VEGFA | Blank | Blank | POS |
| 10 |  |  |  |  |  |  |  |  |  |  |  |  |  |  |

### H. felis infection induces severe inflammation of the gastric corpus

Chronic inflammation associated with *Helicobacter* infection is a primary risk factor for gastric cancer initiation. Our studies demonstrate that HP and HF elicit robust macrophage activation in vitro. Next, we assessed the gastric immune-cell landscape 2 months post-challenge to determine how these bacterial species affect gastric inflammation. Immunostaining of the gastric corpus with the common leukocyte antigen CD45 revealed relatively few immune cells in the mock-challenged stomach, which is consistent with previous reports that the healthy stomach harbors relatively few resident leukocytes (Figure 2A) ^(29, 30)^. HP infection induced modest leukocyte infiltration within the gastric corpus. In contrast, HF infection induced severe leukocyte infiltration throughout the entire gastric corpus (Figure 2A). Next, we utilized spectral flow cytometry to investigate the specific immune-cell populations that respond to *Helicobacter* infection. For these studies, we disassociated the entire gastric corpus. Compared to mock controls, both HP and HF infection induced a statistically significant increase in gastric immune infiltration (Figure 2B). However, the HF-infected stomach had significantly more inflammation than the HP-infected stomach. Both CD3+ T cells and B220+ B cells were significantly increased in HF-infected mice, with T cells being the most abundant immune-cell population (Figure 2B). While T cells were increased in HP-infected stomachs, the increase was not statistically significant. Macrophages were not significantly changed by either *Helicobacter* infection. Neutrophils and eosinophils were only significantly increased in HF-infected mice.

**Figure 2.**
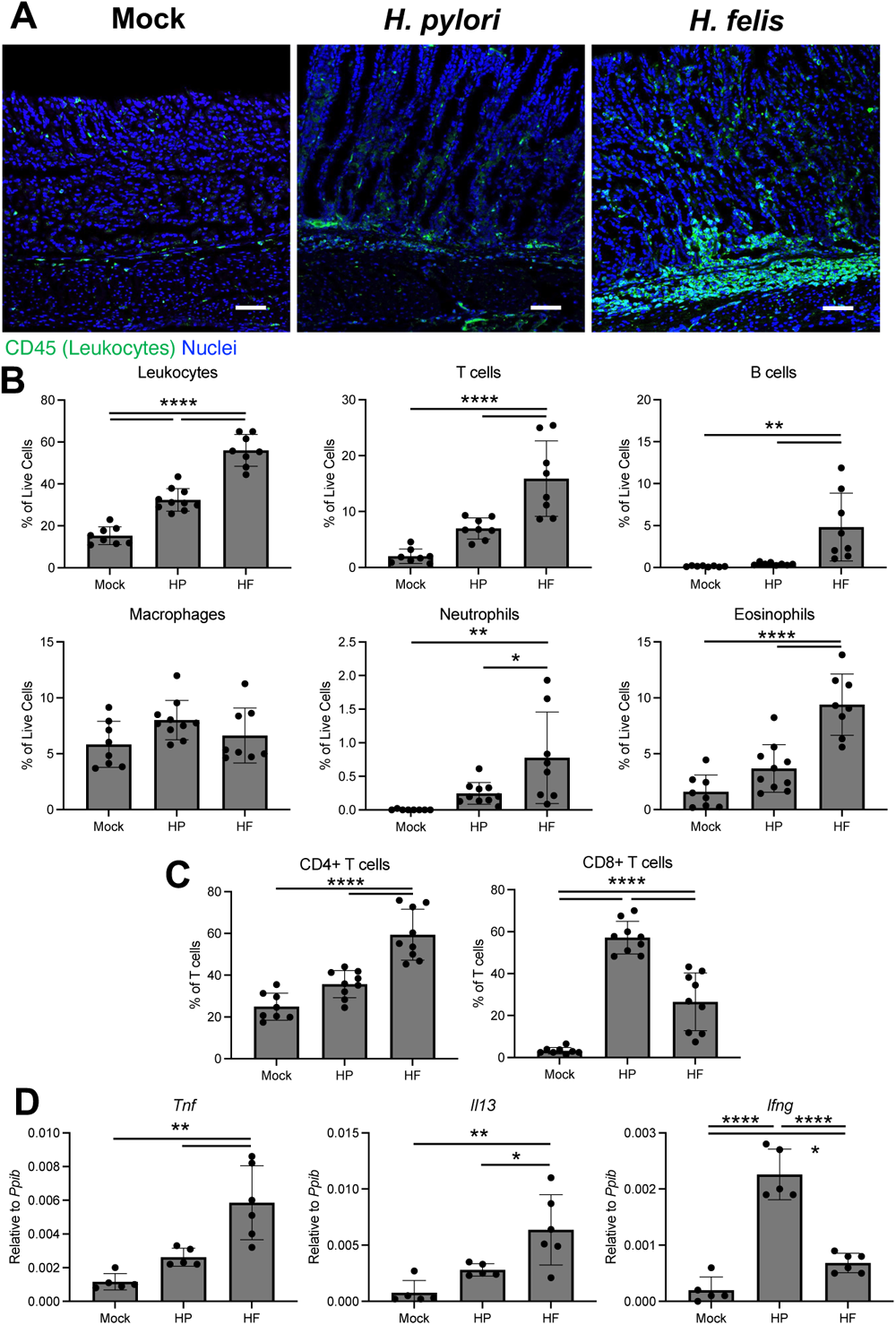
*H. felis-*infected mice develop more extensive gastric inflammation than *H. pylori*. Cryosections (**A**), cells (**B-C**), and RNA (**D**) were collected from the gastric corpus 2 months after mock infection or challenge with *H. pylori* or *H. felis.* (**A**) Representative immunostaining. Sections were probed with antibodies against the common leukocyte antigen CD45 (green). Nuclei were stained with DAPI. n≥6. Scale bars=100µm. (**B-C**) Flow cytometry of the indicated cells types from disassociated gastric corpus. (**C**) CD4+ and CD8+ T cell ratios. n≥8. (**D**) Quantitative RT-PCR of RNA from the gastric corpus for the indicated genes. n≥5. *p≤0.05; **p≤0.01; ***p≤0.001; ****p≤0.0001

Gastric infiltrating T cells are linked to most of the gastric pathologies associated with *Helicobacter* infection in mice ^(8)^. Gastric T cells are rare in the murine stomach during steady-state conditions (Figure 2B), but those T cells present in mock-infected mice were almost entirely CD4+ Helper T cells. Interestingly, while HP induced only a modest increase in total gastric T cells, each bacterial strain differentially affected the CD4/CD8 T cell ratios. HP induced a CD8+ biased T cell response, while HF induced a CD4+ biased T cell response (Figure 2C). Finally, we assessed the expression of inflammatory genes within the gastric corpus by qRT-PCR. *Tnf* and *Il13* transcript levels were not significantly increased in response to HP infection, while both cytokines were significantly increased by HF infection (Figure 2D). In contrast, *Infg* levels were significantly increased in both the HP and HF-infected groups but were highest in HP-infected mice. Together, these results demonstrate that while the HP PMSS1 strain induces gastric inflammation, it is modest when compared to HF.

### H. felis induces more advanced atrophic gastritis than H. pylori in mice

Because HF induced significantly more overall immune cell infiltration and gastric T cell recruitment, we hypothesized that gastric epithelial damage would also be more severe in HF-infected mice. The morphology of the gastric corpus was assessed by H&E. Compared to mock-infected controls, HP-infected mice exhibited a slight increase in inflammatory infiltrate, similar to our findings above in Figure 2. In addition, while the overall gland length appeared unchanged, there was a noticeable foveolar hyperplasia, with the pit cells extending down to the gland neck (Figure 3A). However, the parietal cells and chief cell compartment appeared grossly normal. In stark contrast, HF induced significant changes to the gastric epithelium throughout the entire corpus with a massive increase of immune cells and thickening of the gastric mucosa (Figure 3A). Moreover, there was a dramatic expansion of the gastric pit cells and mucous neck cells, as well as a total absence of parietal and chief cells. Next, we immunostained with the mucous neck cell marker GSII lectin, the mature chief cell marker MIST1 (BHLHA15), and the parietal cell-specific hydrogen-potassium ATPase to better visualize changes to the major gastric cell lineages. HP-infected mice appeared similar to mock-infected controls, with normal proportions of mucous neck, chief, and parietal cells (Figure 3B). Moreover, qRT-PCR for the cell lineage markers *Tff2* (mucous neck cells)*, Gif* (chief cells), and *Atp4b* (parietal cells) demonstrated that the relative expression of these major gastric lineages did not change 2 months post-HP challenge (Figure 3C). In contrast, HP-infected mice exhibited dramatically increased GSII immunostaining, complete absence of MIST1+ cells, and only a few scattered parietal cells (Figure 3B). Similarly, qRT-PCR demonstrated a significant expansion of *Tff2* and a significant 8.9-fold and 4.3-fold loss of *Gif* and *Atp4b,* respectively (Figure 3C). These results demonstrate that HP infection induces non-atrophic gastritis in the gastric corpus within 2 months post-challenge but that HF quickly induces atrophic gastritis denoted by massive damage to the gastric epithelium, leading to dramatic changes to the major gastric epithelial cell lineages.

**Figure 3.**
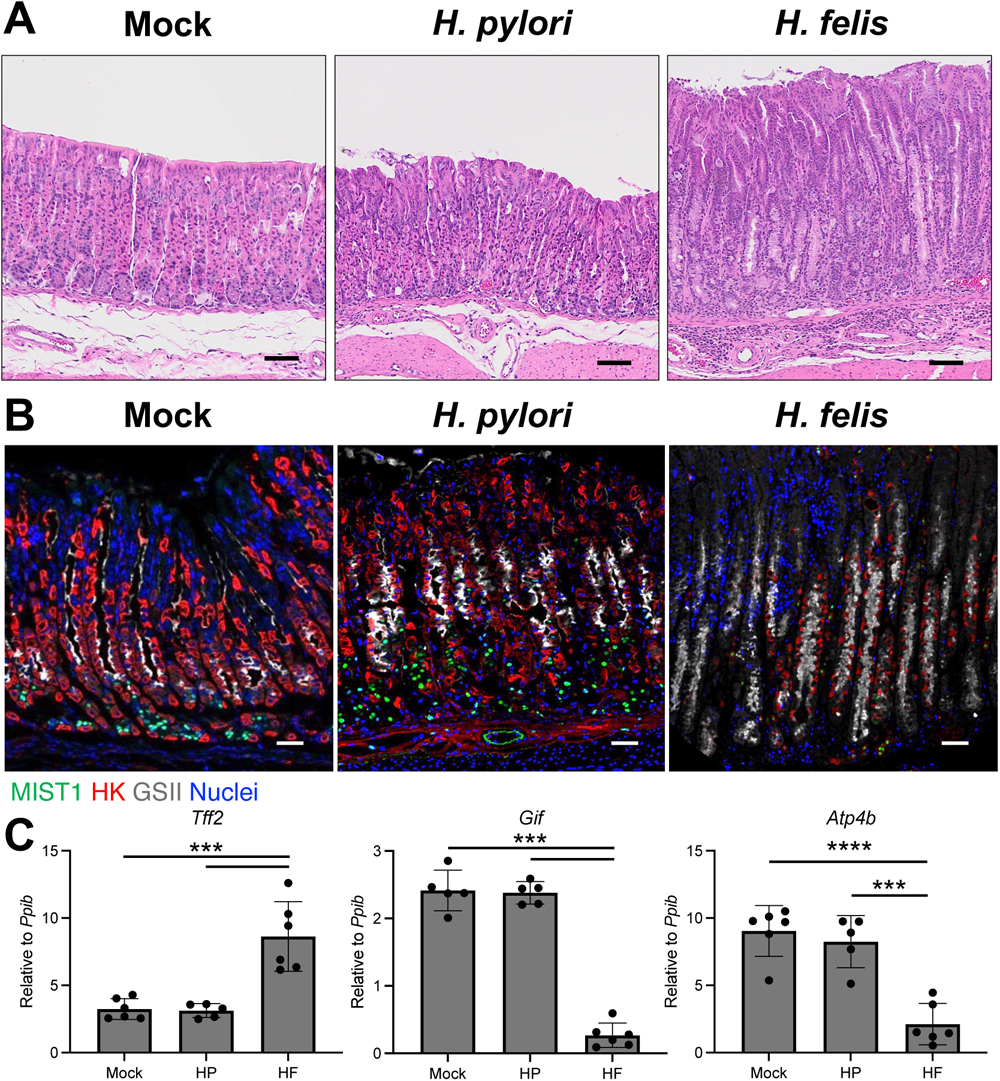
*H. felis-*infected mice develop more extensive atrophic gastritis than *H. pylori*. Tissues (**A-B**) and RNA (**C**) were collected from the gastric corpus 2 months after mock infection or challenge with *H. pylori* or *H. felis.* (**A**) Representative H&E micrographs. (**B**) Representative immunostaining. Sections were probed with antibodies against the MIST1 (mature chief cells, green) the H+/K+ ATPase (parietal cells, red), and the *Griffonia simplicifolia* lectin (mucous neck cells, grey). Nuclei were stained with DAPI. (**A-B**) n≥6. Scale bars=100µM. (**C**) Quantitative RT-PCR of RNA from the gastric corpus for the indicated genes. n≥5. **p≤0.01; ****p≤0.0001.

### H. felis infection induces pyloric metaplasia

The gastric corpus responds to inflammation and glandular damage by modifying cellular differentiation, leading to the development of pyloric metaplasia (PM), a putative preneoplastic lesion ^(31)^. To assess PM development, we immunostained with the *de novo* PM marker CD44v9^(32)^. Mock-infected mice did not exhibit any detectable CD44v9 expression within the glands of the gastric corpus (Figure 4A). Similarly, CD44v9 staining was largely absent within the corpus glands of HP-infected mice, although there were occasionally positive cells within the gland base. In contrast, HF-infected mice exhibited widespread CD44v9-positive glands throughout the gastric corpus glands. Next, we assessed the expression of the PM marker transcripts *Wfdc2* and *Cftr.* Curiously, these transcripts were significantly increased in both HP-and HF-infected mice (Figure 4B), and *Cftr* expression was significantly higher in HP-infected mice compared to HF-infected mice. Overall, HP-infected mice do not have gross morphological features of PM (Figure 4A-B) and lack expression of CD44v9 (Figure 4A) at 2 months post-infection

**Figure 4.**
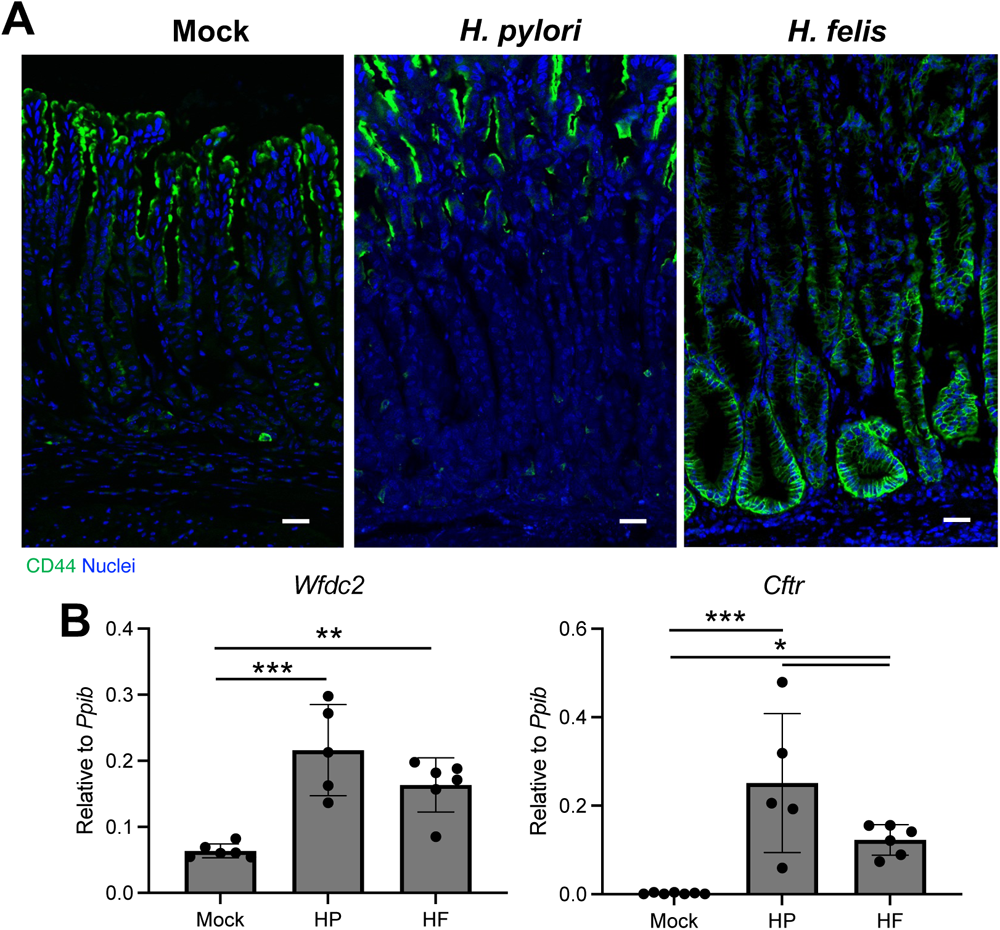
Pyloric metaplasia is more extensive in *H. felis*-infected mice. Tissues (**A**) and RNA (**B**) were collected from the gastric corpus 2 months after mock infection or challenge with *H. pylori* or *H. felis.* (**A**) Immunostaining for the pyloric metaplasia marker CD44v9. n≥5. Scale bar=100µM. (**B**) Quantitative RT-PCR of RNA from the gastric corpus for the indicated genes. n≥6. **p≤0.01; ****p≤0.0001.

### H. felis drive more extensive inflammation in the gastric antrum

While HP is well-adapted to the harsh gastric environment, the extremely low pH of the gastric corpus is toxic and typically inhibits HP colonization, although this colonization pattern is strain specific ^(33)^. As a result, HP usually initially colonizes the gastric antrum and, over time, colonizes the corpus through the lesser curvature ^(31)^. Next, we used H&E micrographs to assess the gross morphology of the gastric antrum in HP and HF-infected mice. Compared to mock-infected controls, the antrum of HP and HF-infected mice developed modest leukocyte infiltration (Figure 5). However, leukocyte infiltration was more extensive in HF-infected mice, and the glands were thickened. These results indicate that HF drives more extensive inflammation throughout the entire stomach, affecting both the corpus and pylorus.

**Figure 5.**
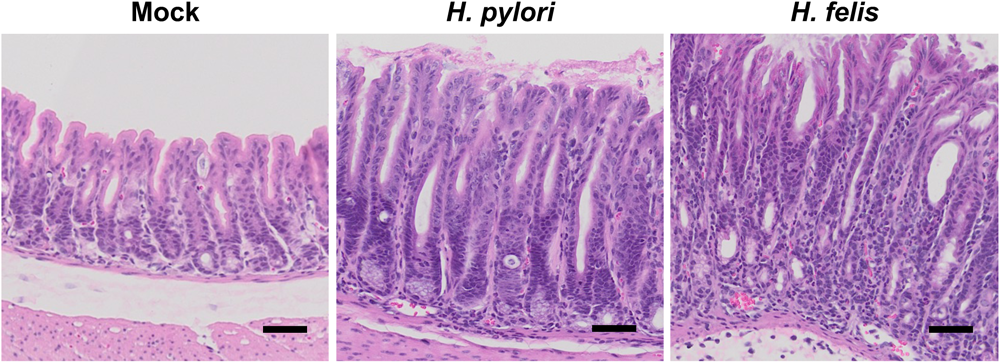
*H. felis* induces more extensive inflammation within the gastric antrum. H&E micrographs of the gastric antrum collected 2 months after mock-infection or infection with *H. pylori* or *H. felis.* Scale bar=50µM. n≥5.

## Discussion

Gastric-colonizing *Helicobacter* species have a complex relationship with the host immune response. Mild inflammation likely benefits the pathogen by causing oxyntic atrophy, allowing further colonization of the gastric corpus ^(31, 34)^ and increasing the expression of adhesion molecules by damaged and metaplastic gastric glands ^(35, 36)^. Severe inflammation is linked to lower bacterial loads due to increased pathogen clearance by the immune system and increased competition due to stomach colonization by intestinal flora as a result of gastric achlorhydria ^(37, 38)^. Severe inflammation ultimately may lead to *Helicobacter* eradication by the host. Therefore, HP has evolved to evade the host immune system ^(27)^. HP LPS lipid A is tetra-acetylated and poorly activates TLR4 ^(24, 39)^. HP flagella mutations at the TLR5 binding site evade TLR5 activation ^(28)^, and HP DNA is modified to avoid TLR9 activation ^(40)^. Rather, HP and HF are potent activators of TLR2 ^(24, 41)^. HF LPS has a similar effect on TLR4 activation, but unlike HP, it is reported to activate TLR9 to induce gastric inflammation ^(42)^. Our RNAseq analysis revealed that HF expressed higher levels of LPS synthesis genes and flagellar genes than HP. Moreover, HF induced significantly more robust inflammatory responses in mice. We, therefore, hypothesized that HF is innately more immunogenic than HP. Surprisingly, while HF induced higher mRNA levels of Il6 and Il1b, both bacterial species induced similar cytokine responses when assessed by a semi-quantitative dot plot array. Similar findings were reported by Mandell *et al.*, where both HP SS1 and HF elicited similar activation of TLR4 and TRL2 ^(24)^. These data suggest that the enhanced inflammation observed in HF-colonized mouse stomachs is not due to enhanced immunogenicity of HF and may result from other mechanisms such as enhanced HF colonization or HP-mediated suppression of inflammatory pathways.

Our mouse colonization studies demonstrated that HF induced significantly more extensive gastric inflammation and epithelial remodeling than HF. Chronic inflammation is necessary and sufficient for gastric cancer initiation. Mice that lack mature B and T cells are protected from morphological changes to the gastric epithelium in response to infection by either HP or HF ^(8, 43)^, and myeloid cell activation is required for pyloric metaplasia initiation ^(29, 30, 44)^. Moreover, overexpression of the proinflammatory IL1B or IFN-gamma under control of the parietal cell-specific H+/K+ ATPase β promoter drives gastric neoplasia development without accompanying *Helicobacter* infection ^(4, 5)^. While we found that both HP and HF induced similar macrophage activation in vitro, gastric inflammation 2 months post-infection was significantly different for these two bacterial species. HP elicited only mild inflammatory responses 2 months post-infection. In contrast, HF induced severe inflammation throughout the gastric corpus at the same time point. While previous studies reported that HP triggered gastric inflammation, atrophy, and metaplasia by 2 months post-infection ^(16, 45, 46)^, our findings indicate that these events are mild and restricted to the gastric corpus lesser curvature. HP typically initially colonizes the gastric antrum and, over time, spreads through the corpus lesser curvature ^(47)^. While the underlying mechanisms of this pattern of spread through the stomach is not fully defined, the lesser curvature is more susceptible to inflammation and consequential oxyntic atrophy ^(31, 44)^. In contrast, HF-induced metaplasia and inflammation were ubiquitous throughout the gastric corpus. Within the gastric antrum, both bacterial species induced inflammation but as within the corpus, HF triggered more extensive inflammation. Poor colonization by the PMSS1 strain may account for some of the poor inflammation. Sigal *et al*. found that HP PMSS1 is present in the corpus but does not colonize gastric corpus glands of C57BL/6J mice 2 months post-challenge ^(26)^. Moreover, interactions with the microbiota may abrogate HP-induced inflammation as the same study reported that C57BL/6 mice from different vendors exhibit dramatic differences in corpus inflammation ^(26)^. These factors do not appear to affect HF to the same extent, as the reported gastric phenotypes associated with HF have remained largely consistent over 2.5 decades ^(8, 20, 42, 48-50)^.

T cells are required for the bulk of gastric epithelial damage and remodeling associated with *Helicobacter* infection in mice ^(8, 51)^. Elevated levels of Th1/Th17-associated cytokines are linked to increased cancer risk in humans ^(52, 53)^. Here, we found that HP does not induce a significant T cell response 2 months post-infection. This lack of response is consistent with the mild epithelial changes and lack of metaplasia development. HF induced dramatic T cell activation, particularly in the CD4+ T cell compartment. CD4+ T cells are reported to drive the bulk of gastric damage associated with HF infection ^(54)^. Interestingly, while HP only caused mild gastric T cell recruitment, it caused a shift in the CD4/CD8 T cell ratio, skewing towards an enhanced CD8 cytotoxic T cell response. This increase is likely due to the CagA pathogenicity island and functional T4SS that transfers bacterial components into host epithelial cells ^(55)^. We found that while HF inflammation was predominated biased towards CD4+ T cells there still was an overall increase in gastric CD8+ T cells. CD8 T cells have also been reported to induce gastric damage during HF infection, but their role is secondary to CD4 T cells ^(49)^. The exact role of these cytotoxic T cells is unclear. While CD8+ T cell responses contribute to epithelial damage ^(49, 55)^, they undoubtedly also play important roles in responding to neoantigens to remove developing tumor cells. The hyper-inflamed environment induced by *Helicobacter* may promote cytotoxic T cell exhaustion and dysfunction, potentially explaining why more intense gastric inflammation is linked to increased cancer risk.

HP and HF are closely related *Helicobacter* species, sharing more than 60% of each other’s genome ^(19)^. HF’s genome encodes many HP orthologs that are required for gastric colonization, such as a urease cluster, flagellar and chemotactic genes, gastric epithelial adhesion proteins, RecA, collagenase, and iron uptake gene clusters. CagA and the vacuolating toxin VacA are the most well-known HP virulence factors associated with gastric cancer risk in humans and are notably absent from the HF genome ^(19)^. CagA is a bonafide bacterial oncoprotein that is injected into host epithelial cells through its T4SS ^(56)^. Once inside host cells, CagA attaches to the plasma membrane and promiscuously interacts with host proteins, inducing mutations, cellular reprogramming, and inflammation ^(57)^. Transgenic mice that ectopically express CagA from parietal cell-specific H+/K+-ATPase promotor spontaneously develop gastric tumors ^(58)^. While CagA is sufficient to drive gastric inflammation and cancer initiation, our results indicate that inflammation induced by CagA+ HP still pales in comparison to HF. CagA expression is likely one of the most important rationales for using HP in mice infection studies. However, when HP was re-isolated from infected mice, it was found that they had lost their T4SS functionality by 3-4 months post-infection ^(16, 17)^. Thus, the selection of HP for mouse infection studies strains solely on their expression of CagA may not yield the desired phenotypes.

In summary, HP infection studies in mice mount specific challenges in modeling the inflammatory phenotypes and cancer initiation observed in humans. These problems stem from the fact that HP is not a natural mouse pathogen and common mouse-adapted strains such as PMSS1 suffer from weak immunogenicity and functional loss of virulence factors that promote cancer risk. HF has been used for decades to model pro-neoplastic gastric inflammation and cancer initiation and quickly and consistently initiates inflammation in mouse models. While HF is more strongly suited for studies on pathogenesis and cancer initiation, its lack of CagA and VacA limits its utility for studies on the direct effects of the bacteria on gastric epithelial cells. Moreover, HP is likely better suited for studies on bacterial colonization and interactions with the gastric microbiota.

## Acknowledgments

This work was supported by West Virginia University start-up funds (J.T.B) and a grant from National Institutes of Health P20GM121322 (J.T.B.). The West Virginia University Microscope Imaging Facility and Flow Cytometry & Single Cell Core receive support from the National Institutes of Health grants P30GM103503 and S10 grant OD028605, respectively. The authors thank Richard Peek Jr. (Vanderbilt University Medical Center) for technical assistance.

